# 3D Microphysiological Tumor Model for Dual-Targeting CAR T Cell Immunotherapy

**DOI:** 10.1101/2024.08.16.608227

**Authors:** Xuan Peng, Željko Janićijević, Liliana Rodrigues Loureiro, Lydia Hoffmann, Poh Soo Lee, Isli Cela, Benjamin Kruppke, Alexandra Kegler, Anja Feldmann, Michael Bachmann, Larysa Baraban

**Affiliations:** Helmholtz-Zentrum Dresden-Rossendorf, Institute of Radiopharmaceutical Cancer Research, 01328 Dresden, Germany; Faculty of Medicine and University Hospital Carl Gustav Carus, Technische Universität Dresden, 01307 Dresden, Germany; Max Bergmann Center of Biomaterials and Institute of Materials Science, Technische Universität Dresden, 01069 Dresden, Germany; National Center for Tumor Diseases (NCT), Dresden, Germany; German Cancer Research Center (DKFZ), Heidelberg, Germany; German Cancer Consortium (DKTK), Dresden, Germany

**Keywords:** droplet microfluidics, PEGDA hydrogel beads, prostate cancer, fibrosarcoma, immunotherapy, solid tumor, tumor microenvironment, fibroblast activation protein, immunostaining

## Abstract

The efficiency of immunotherapy stays limited for solid tumors. It is mainly caused by the tumoral structural heterogeneity and its complex microenvironment, which impede the infiltration of immune cells into malignant tissues. Mimicking this environment in frames of microphysiological models remains a challenge, significantly increasing costs of the clinical translation for the new therapies. Here, we study a 3D multi-spheroid model incorporating prostate stem cell antigen (PSCA) modified PC3 human prostate cancer cells and fibroblast activation protein (FAP) expressing fibrosarcoma HT1080 cells embedded within the soft hydrogel microbeads. We use this model to trial the immunotherapy based on the universal chimeric antigen receptor (UniCAR) T cells, and to better understand the impact of FAP on the immunotherapeutic treatment of solid tumors. First, we demonstrate the successful chemoattraction and infiltration of UniCAR T cells into the area of solid tumors, as well as the ability of UniCAR T cells to navigate through artificial extracellular matrix barriers. We further observe the synergistic efficacy of a dual-targeting UniCAR T cell approach against FAP and PSCA antigens, which represent the tumor microenvironment and the tumor, respectively. The results of our studies offer valuable methodologies and insights for engineering different 3D tumor models and studying immunotargeting of small-sized solid tumors (e.g., metastases and residual tumors). The developed microphysiological system has great potential to advance cancer research efforts aiming to elucidate the pivotal role of microenvironment in solid tumor development, enabling therapy trials and more precise prognosis for patients.

## 1. Introduction

The complexity of cancer and its protective mechanisms limits the therapeutic efficiency of cancer immunotherapy to around 30-40% in clinical patients.^1, 2^ The tumor microenvironment (TME) is deemed responsible for the lower efficacy of immunotherapy in solid tumors compared to blood-related malignancies.^1^ Rapid translation of effective treatments remains hindered by the lack of suitable cancer models recapitulating the inherent tumor heterogeneity, resistance to therapy, disease progression, and patient-to-patient variability.^3^ Numerous efforts have been dedicated to preclinical cancer modeling,^4, 5^ particularly driven by the advances in materials science and three-dimensional (3D) cell culture technologies.^6–8^ Various biomaterials ranging from biologically derived polymers (such as matrigel^9^, collagen^10^ and alginate^11–13^) to synthetic polymers (such as poly(ethylene glycol) diacrylate (PEGDA)^14–16^ and carboxymethyl cellulose^17^) have been studied to integrate the complex cellular assemblies, while only a minority of current 3D cancer models could be reconstructed in the existing biomaterial-based matrices.^3^ Spheroids, organoids, microfluidic systems, and bioprinting solutions have emerged as representative platforms for improved recapitulation of the physiological aspects of human cancers and their intricate interactions with the TME.^6, 18–22^ However, accurate replication of the complex and dynamic nature of the TME, at both, structural and biochemical, signaling levels remains a significant challenge.^23–25^ The mimicking of key TME properties is still challenging primarily due to: (1) the insufficient understanding of the role of physical cues (e.g., stiffness, viscosity, and viscoelasticity) that regulate tumor growth and invasion;^26–28^ (2) the impact of biomolecules that remodel the TME, influence the metastatic ability of cancer cells, and modulate immunotherapeutic responses; ^29, 30^ (3) the stromal cells (e.g., cancer-associated fibroblasts (CAFs) and adipocytes) which play crucial roles in tumor metabolism, growth, and metastasis;^31^ (4) the distinct properties of certain cancer cell types, such as poor aggregation and limited stability in culture; For instance, human prostate cancer cell (PC3) spheroids, reveal loose aggregation and short-term survival for a maximum of *c.a.* ten days.^32, 33^

To the best of our knowledge, most of the current 3D tumor models are focused on increasing biodiversity and mimicking tumor structures. However, biomolecules, such as fibroblast activation protein (FAP) which contributes to tumor progression and offers a promising target for cancer therapy,^34–36^ were rarely considered when building a 3D tumor model. The reasons described above restrict the possibilities of reaching sufficient spheroid maturation for use in immunotherapy (incl. chimeric antigen receptor (CAR) T cell therapy) investigations.^37, 38^ Thus, it is imperative to develop more adaptable 3D tumor models that offer the possibility to tune both, physical (e.g., mechanical, Figure 1a(I)) and chemical, properties of the microenvironment, as well as to demonstrate novel therapeutic approaches *in vitro*.

**Figure 1.**
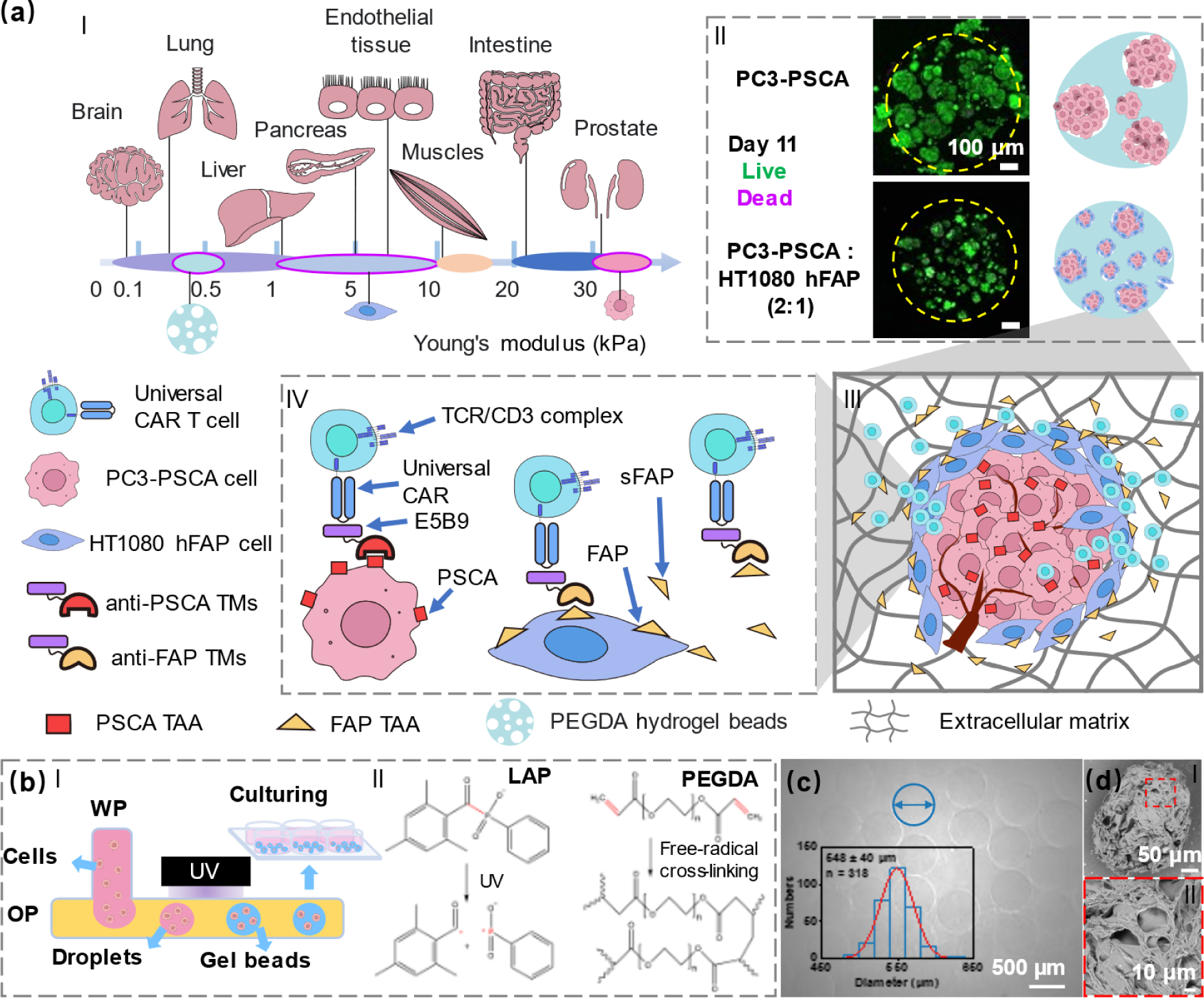
Conceptual illustration of the 3D tumor model and PEGDA hydrogel beads generation. (a) Conceptual illustration of tumor tissue engineering for investigating UniCAR T cell therapy. As reported, the stiffness of TME is important for engineering 3D tumor models, and we considered Younǵs modulus of main human organs in our design (Ⅰ).^46^ In this study, the 3D tumor models were created using PSCA-modified PC3 human prostate cancer cells (PC3-PSCA) and FAP-modified HT1080 fibrosarcoma cells (HT1080 hFAP) embedded in PEGDA hydrogel beads of adjustable stiffness, showing good viability and different morphologies after 1 week of culturing (Ⅱ). We apply this construct to model some key parameters within the TME including cancer cells, stromal cells, extracellular matrix, and soluble factors (Ⅲ).^10^ A pilot UniCAR T cell therapy study was conducted using anti-PSCA and anti-FAP target modules (TMs) to target the corresponding tumor-associated antigens (TAA) (Ⅳ). (b) Schematic illustration of the T junction-based microfluidic system coupled with the UV light source for the fabrication of PEGDA hydrogel microbeads (Ⅰ); Mechanism of the PEGDA photopolymerization process (Ⅱ).^14, 47^ (c) A representative image of PEGDA hydrogel beads generated under the average UV intensity of 0.23 W‧cm^−2^; The used flow rate for water phase (WP) was 5.5 μL‧min^−1^ and for oil phase (OP) 110 μL‧min^−1^ (inset shows the distribution of generated PEGDA bead diameters indicating high monodispersity). (d) Representative SEM images of freeze-dried PEGDA hydrogel beads showcasing the presence of macroscopic pores in the dry state. PSCA: prostate stem cell antigen, FAP: fibroblast activation protein, sFAP: soluble fibroblast activation protein. TMs: target modules, PEGDA: poly(ethylene glycol) diacrylate, LAP: lithium phenyl-2,4,6-trimethylbenzoylphosphinate.

In recent decades, immunotherapy and the treatment with CAR T cells, in particular, has become a central focus for engaging the immune system in the fight against cancer, producing remarkable preclinical and clinical progress.^39–41^ To make CAR T cell therapy safer and more controllable, modular approaches are under active development, such as the so-called universal CAR (UniCAR) approach.^42, 43^ A key component of the UniCAR T cell system is the target module (TM), a compound aiming to mediate the linkage between effector and target cells (Figure 1a(IV)). By tuning the types and concentrations of specific TMs, it is possible to guide and redirect UniCAR T cells in an individualized time– and target-dependent manner.^44^ TMs include two components: (1) a peptide epitope against a UniCAR-engineered T cell, and (2) a binding moiety against the specific tumor-associated antigen.^45^ UniCAR T cells can bind reversibly to single or multiple TMs. Due to the complex nature of CAR T cell immunotherapy and potential side effects, the development of reliable 3D *in vitro* tumor models is a promising approach to enable systematic investigations and acquire a deeper understanding of interactions between CAR T cells and targeted tumors in the microphysiological systems.

In this study, we aim to contribute to the research about unlocking the tumor microenvironment for immunotherapeutic treatment and construct the microphysiological 3D multi-spheroid tumor model in hydrogel microbeads, using PSCA-modified prostate cancer cells (PC3-PSCA) cocultured with the HT1080 (HT1080 hFAP) cells as a source of FAP in the microenvironment. We use this model, to investigate the immunotherapeutic efficiency of UniCAR T cells in the presence of both, anti-PSCA and anti-FAP directed TMs (Figure 1a (Ⅲ&Ⅳ)). PEGDA hydrogel beads with an optimized elastic modulus are used here to mimic the native organ and extracellular matrix stiffness (Figure 1a (Ⅰ&Ⅱ)), enabling miniature, transparent, and reproducible microenvironments to facilitate long-term cell culturing (up to 3-4 weeks), immunohistochemistry staining, and T cell monitoring. The results of our investigations based on the constructed microphysiological 3D tumor model indicate: (1) pronounced differences in the morphology of tumor structure with and without FAP presence in the environment; (2) the ability of UniCAR T cells to efficiently infiltrate the tumor region and penetrate artificial extracellular matrix barriers; (3) inhibiting role of FAP or soluble FAP (sFAP) in the therapeutic activity of UniCAR T cells; and (4) the synergistic effect on cancer cell killing in a dual-targeting UniCAR T cell therapy. This study holds promise for the facile design and engineering of *in vitro* 3D tumor models with tunable physical and biochemical elements of complex TME, showcasing a potential route to expedite the translation of CAR T cells into clinical immunotherapy procedures. Moreover, it provides valuable methodologies and insights for modeling therapies of small metastatic or residual tumors that cannot be easily resolved with traditional medical imaging modalities, and thus cannot be timely treated.

## 2. Results

### 2.1 Structure and mechanical properties of PEGDA hydrogel beads

To generate PEGDA hydrogel beads, we designed a droplet-based microfluidic platform coupled to a 365-nm UV lamp for photopolymerization (Figures 1b (Ⅰ) and S1 a). Initially, the water phase (WP) containing 10% (w/v) of PEGDA monomers and 1% (w/v) of lithium phenyl-2,4,6-trimethylbenzoylphosphinate (LAP) photoinitiator dissolved in cell culture medium, and the mineral oil phase (OP), were separately injected into fluorinated ethylene propylene (FEP) tubes (outside diameter: 1.6 mm; inside diameter: 0.5 mm) with controlled flow rates using syringe pumps (Cetoni 210 base, q_WP_ = 5.5 μL‧min^−1^, q_OP_ = 110 μL‧ min^−1^). The T-junction facilitated the production of reproducible and monodisperse water-in-oil emulsion droplets that were further rapidly polymerized using UV irradiation (mean intensity: 0.23 W‧cm^−2^; exposure time: 2.5 s), resulting in hydrogel beads with the average diameter of 548±40 µm (Figure 1c). The measurements and calculations of UV irradiation parameters are explained in sections 4.2 and SI, as well as in Figure S1b,c. The polymerization reaction was achieved through the efficient absorption of light at 365 nm by the LAP photoinitiator, which resulted in the production of free radicals initiating the polymerization process (Figure 1b (Ⅱ)),^14, 47^ entailing chain growth proceeding via covalent cross-linking through the carbon double bonds of acrylate side groups terminating the PEGDA monomers.^15^ Cross-linked PEGDA networks form hydrogels in aqueous media that, in our case, assume the shape of spherical beads with high monodispersity.

Fourier-transform infrared spectroscopy analysis confirmed the successful formation of the cross-linked PEGDA polymer network,^15, 48^ indicating no significant influence of the complex RPMI STINO cell culture medium on the final chemical composition of the hydrogel (Figure S2, SII Note). The hydrogel beads represent a scaffold formed as a porous 3D cross-linked polymer network (Figure 1d), providing good mechanical stability as well as sufficient space for cell culturing. Prostate cancer cells are sensitive to the mechanical properties of their microenvironment, and the stiffness of hydrogel beads plays a crucial role in cell behavior. Mechanical analysis (MicroTester, Figure S3, see section 4.4 for details) shows that the hydrogel beads exhibit Younǵs moduli ranging from 300 to 600 Pa, increasing with the UV intensity used for bead generation (mean intensity: 0.23-0.35 W‧cm^−2^). The measured stiffness range covers stiffness values obtained for some major organs, such as the liver and lungs (Figure 1a),^49^ representing potential targets for prostate cancer metastasis.

### 2.2 Spheroid culturing within PEGDA hydrogel beads

Both, mono-cultured (PC3-PSCA or HT1080 hFAP) and co-cultured spheroids, where PC3-PSCA and HT1080 hFAP were mixed at different ratios (P:H = 10:1, 2:1), exhibited a robust growth in PEGDA hydrogel beads, sustaining culture for over 20 days. All spheroids exhibited solid cell aggregations. PC3-PSCA and P+H spheroids (with PC3-PSCA to HT1080 hFAP ratios of 2:1 and 10:1), with the same initial total cell numbers, displayed similar growth trends within the first week (Figures 2a and S4). Thereafter, PC3 spheroids showed a faster increase in size compared to P+H spheroids. After two weeks, the average diameter of PC3-PSCA spheroids reached 125 μm, compared to only 50 μm reached by P+H spheroids (for both P:H ratios: 2:1 and 10:1; Figure S5). The smallest HT1080 hFAP spheroids had a highly heterogeneous morphology, making reliable size measurements difficult. Notably, all spheroid types maintained high viability over the culturing period of approximately 20 days (Figures 2b and S6). The diameters of some PC3-PSCA spheroids reached *c.a.* 300 μm, while the diameters of the P+H spheroids (with PC3-PSCA to HT1080 hFAP ratios of 2:1 and 10:1) were rarely larger than 100 μm. The HT1080 hFAP cells alone did not aggregate into regular spheroids. Spheroids tend to reach much smaller sizes for higher proportions of HT1080 hFAP cells in the initial cell culture, indicating that these cells hinder cell-cell aggregation. Such an effect could be attributed to the potential degradation of extracellular matrix components by FAP/sFAP which is expressed by HT1080 hFAP cells.^29, 50^

**Figure 2.**
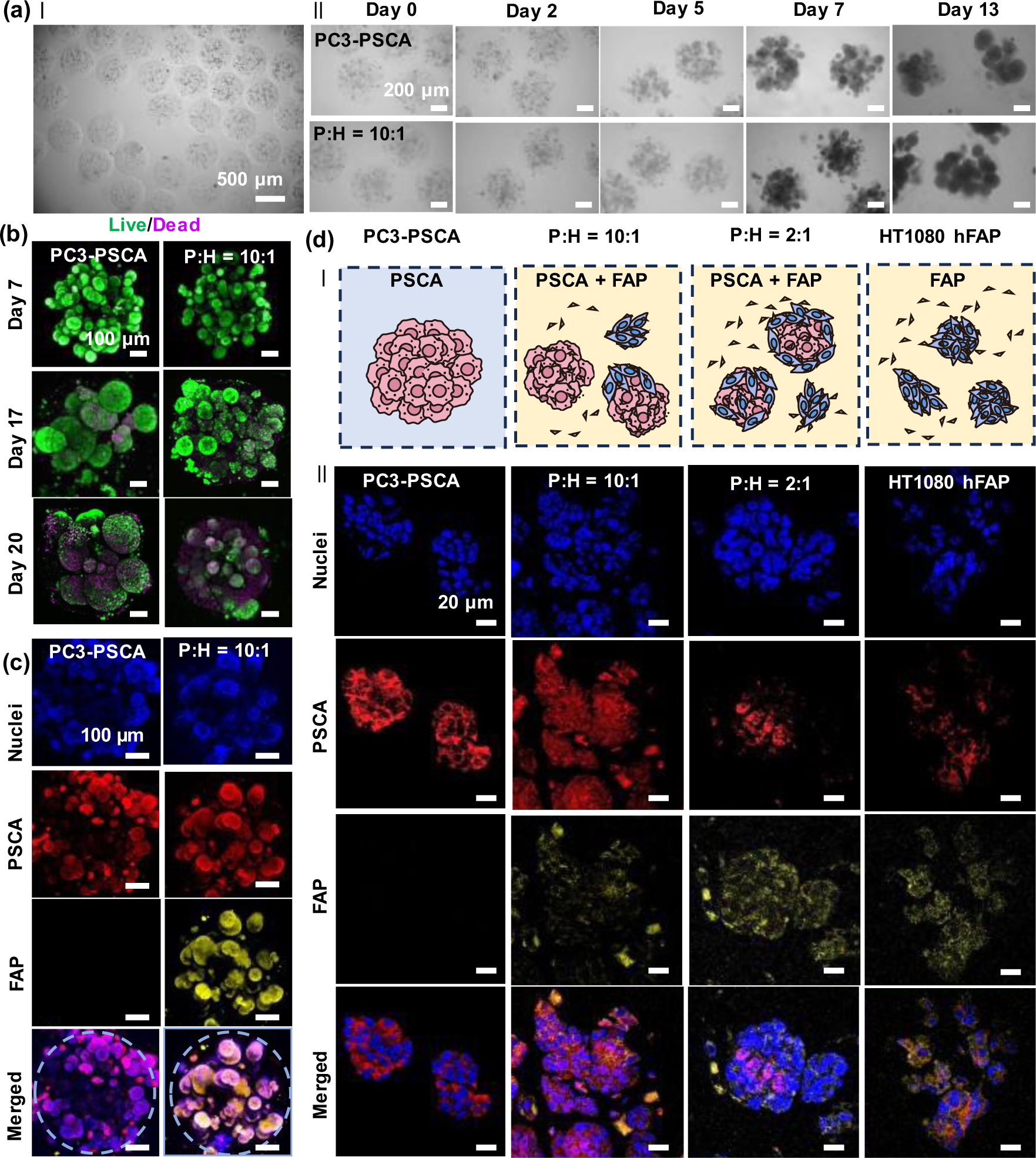
Different spheroids formed in PEGDA hydrogel beads. (a) Representative images showing spheroid formation and proliferation in PEGDA hydrogel beads with PC3-PSCA cells or PC3-PSCA and HT1080 hFAP cells (P+H (10:1): PC3-PSCA to HT1080 hFAP ratio = 10:1). (b) Representative Live/Dead (green/magenta) staining images of PC3-PSCA and P+H spheroids in PEGDA hydrogel beads. (c) Immunostaining images of the structures of PC3-PSCA and P+H spheroids in intact PEGDA hydrogel beads. (d) Illustrations of different spheroid compositions and immunostaining images of spheroid sections on day 8 (cyan: nuclei; red: PSCA; yellow: FAP).

To gain insight into the structure of different spheroids, we conducted fluorescent immunostaining. PC3 cells abundantly express PSCA, while HT1080 cells intensely express FAP (Figure 2d). A lower level of PSCA was also observed in HT1080 hFAP cells (Figure 2d) lacking detectable PSCA on their surface (data not shown), following previous reports.^51^ FAP was observed exclusively on the surface of co-cultured spheroids (Figure 2c). Immunostaining results for sections highlighted the heterogeneous nature of P+H spheroids and the presence of FAP or sFAP (see Figure 2d and S7). HT1080 hFAP was predominant in P+H (2:1) spheroids due to a higher abundance of HT1080 hFAP cells on the surface of spheroids compared to their interior. Interestingly, even when HT1080 hFAP accounted for only 10% of the initial culture, a significant amount of FAP was observed in the P+H (10:1) culturing experiments, thereby eventually indicating the formation of a ‘protective’ layer for the tumor. Thus, our findings indicate that PEGDA hydrogel beads hosting multiple P+H spheroids offer the model, where hFAP cells appear as the stable source of cell-associated and microenvironmental FAP/sFAP, peculiar also to the *in vivo* tumors.^34–36^ This tumor model allows studying CAR T cell therapy within a microenvironment characterized by physical barriers and FAP-mediated immune suppression.

### 2.3 CAR T cell migration and infiltration

#### 2.3.1 T cell migration reproduces *in vivo* behavior

In the following, we performed experimental trials aiming to model the immunotherapeutic effect of UniCAR T cells targeting PSCA and FAP, both expressed in P+H spheroids. Observation of the activation, migration, and infiltration ability of T cells in the complex microenvironment, created by the elastic hydrogel matrix and FAP-positive cells around PC3, is crucial for unlocking the mechanisms of the tumor microenvironment. P+H (10:1) spheroids were selected for UniCAR T cell migration assessment due to their resemblance to both, solid and metastatic, tumors in terms of structure and distribution.^52^ To evaluate the influence of anti-PSCA and anti-FAP TMs in the cell culture medium, the migration statistics of UniCAR T cells in the presence of anti-PSCA and anti-FAP TMs (P+F), anti-PSCA TMs (PSCA), anti-FAP TMs (FAP), and without any TM were compared during the first 60 min of the experiment. UniCAR T cell migration was tracked in the time-lapse movies (Sup_video1) on the well slide surface, within the gel, or on the surface of the spheroids (Figure 3, see section 4.10 for details). The migration statistics of wild-type (WT) T cells were also calculated and used as a control. Interestingly, the displacement distribution images (Figure 3a(Ⅰ) and S8) show that the highest numbers of UniCAR T cells gather around hydrogel beads when both anti-PSCA and anti-FAP TMs are present. To quantify the differences in cell numbers, T cells were counted in a ring area with an inner diameter of 450 μm and an outer diameter of 800 μm (Figure 3b). The number of T cells in the ring area reached a peak after approximately 30 min. In the P+F group, there were approximately 250 more UniCAR T cells on average compared to the starting time point, indicating a higher accumulation trend compared to other groups. In addition, the recorded T cell velocities (Figures 3a(Ⅱ) and S9) indicated the alternating periods of motility and arrest phases of T cells, in accordance with previous reports.^53^ We observed a trend suggesting lower average velocities of T cells in FAP and P+F groups in comparison with those in WT and PSCA groups (Figure S10). The intensified accumulation and lower velocities of T cells in FAP and P+F groups could be attributed to the presence of HT1080 hFAP cells and sFAP in hydrogel beads,^54, 55^ which may synergistically attract UniCAR T cells upon the addition of anti-FAP TMs.

**Figure 3.**
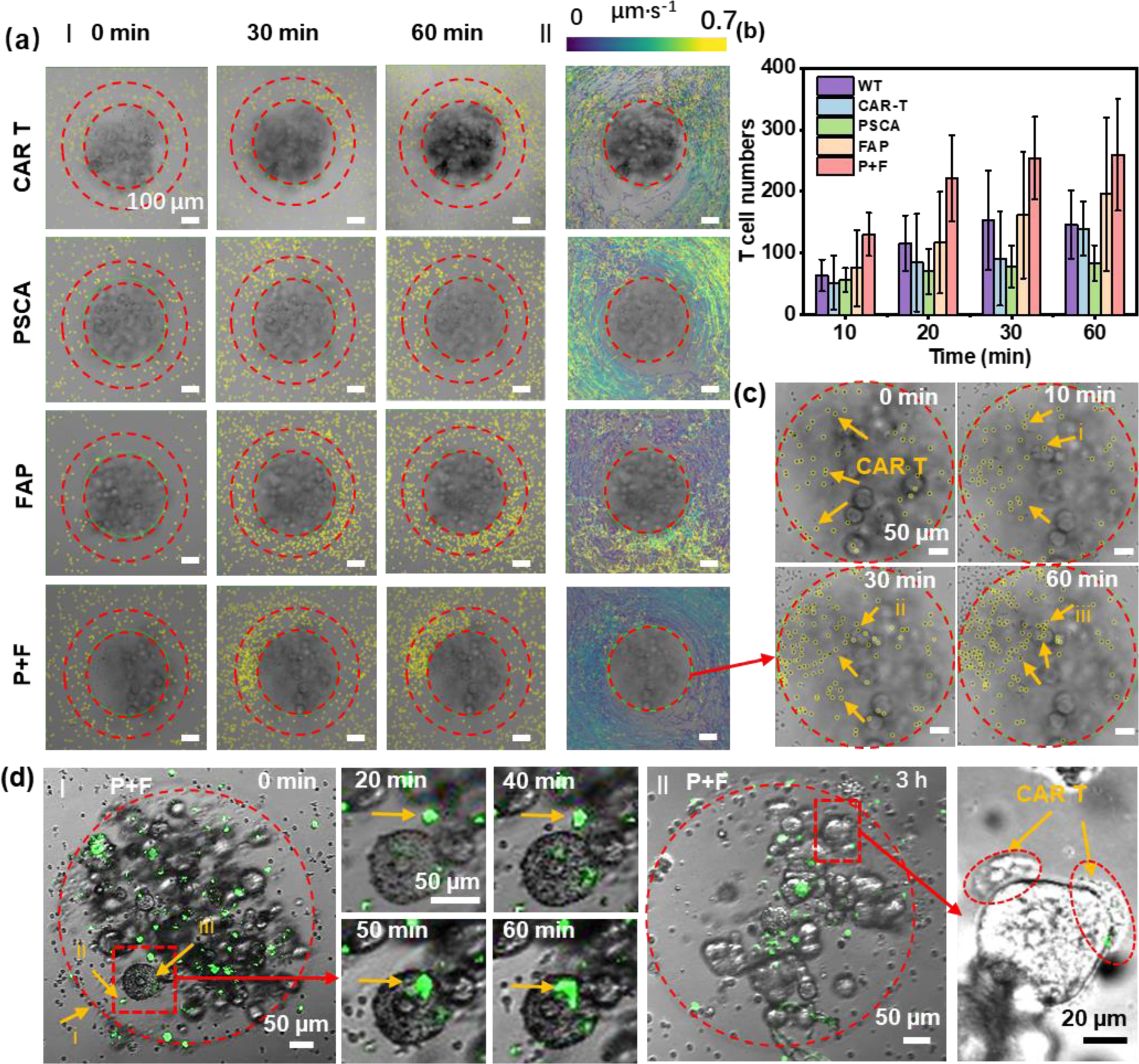
T cell migration study in PEGDA hydrogel beads with P+H (10:1) spheroids. (a) Representative images showing T cell distribution (small yellow circles) in wells under different culturing conditions (I) and tracking of T cell paths over 1 h (II). Each line represents an individual cell track. The color map indicates the instantaneous velocity values for T cells. CAR T: P+H spheroids with UniCAR T cells; PSCA: P+H spheroids with UniCAR T cells and anti-PSCA TMs; FAP: P+H spheroids with UniCAR T cells and anti-FAP TMs; P+F: P+H spheroids with UniCAR T cells, anti-PSCA TMs, and anti-FAP TMs. (b) Average cell counts in ring areas designated in a(I) after culturing the spheroids and T cells for a certain period; the ring area has an inner diameter of 450 μm and an outer diameter of 800 μm, n = 3. (c) Representative images of P+F group from Figure 3a(II) showing T cell distribution inside the hydrogel bead. (d) (I) Representative time sequence of fluorescent micrographs illustrating the T cells approaching and contacting the spheroid within a PEGDA hydrogel bead; (II) Representative images of coordinated T cell accumulation on the surface of a spheroid after 3 h of culturing.

#### 2.3.2 UniCAR T cell accumulation is enhanced by cell-to-cell signaling

UniCAR T cells infiltrated into PEGDA hydrogel beads and reached the spheroids (Figure 3c,d). The cells explored the surface of spheroids during the next few hours (Sup_video2). After around 3 h, distinct regions of T cell accumulation were observed on the surfaces of P+H spheroids cultured with anti-PSCA and anti-FAP TMs, while no clear accumulation regions were detected for other groups (Figure 3a(II) and S8). Coordinated cell gathering at a defined location suggests that the accumulation of UniCAR T cells on the P+H spheroids is not only improved by the dual targeting of PSCA and FAP but also enhanced by cell-to-cell signaling, causing a positive feedback loop. A similar phenomenon was also observed in a 2D plate experiment (Figure S11), confirming recently published observations.^53^ In further text, we go beyond the mere accumulation of UniCAR T cells on the spheroid surface and explore their capability to effectively eliminate cancer cells.

### 2.4 Modeling of immunotherapeutic efficiency

PC3-PSCA spheroids were used as a control to compare the dual targeting efficiency. After 48 h of incubation under different culturing conditions, T cell accumulation (green) and efficient killing (magenta) were observed in PC3-PSCA spheroids cultured with UniCAR T cells and anti-PSCA TMs (Figures S13 and S14). However, the same effects were not evident in P+H spheroids cultured solely with anti-PSCA TMs (Figure 4a,b). Distinct dense accumulations of UniCAR T cells and efficient cell killing were only observed when P+H (10:1) spheroids were cultured with both, PSCA and FAP TMs. Immunostaining of small nuclei and PD-1 confirmed the presence of UniCAR T cells inside hydrogel beads and these cells produced granzyme B to effectively kill tumor cells (Figure 4c). The effective cross-linking between a UniCAR T cell and a cancer cell through TMs can trigger the production of cytokines, including TNF-α and IFN-γ, that are released to the tumor microenvironment (Figure 4a and S12). The lack of significant difference between the PSCA and P+H groups, as observed *in vivo* in a previous report,^56^ could be attributed to the blocking of FAP on the cytokine receptors on the cancer cell membrane. Figure S15 illustrates our hypothesis on the mechanism of the dual-targeting system. A lower cell killing efficiency was observed in P+H (2:1) and HT1080 hFAP groups with a large portion of FAP (Figure S16), indicating that our 3D cancer models are promising for mimicking the TME under different cancer development conditions.

**Figure 4.**
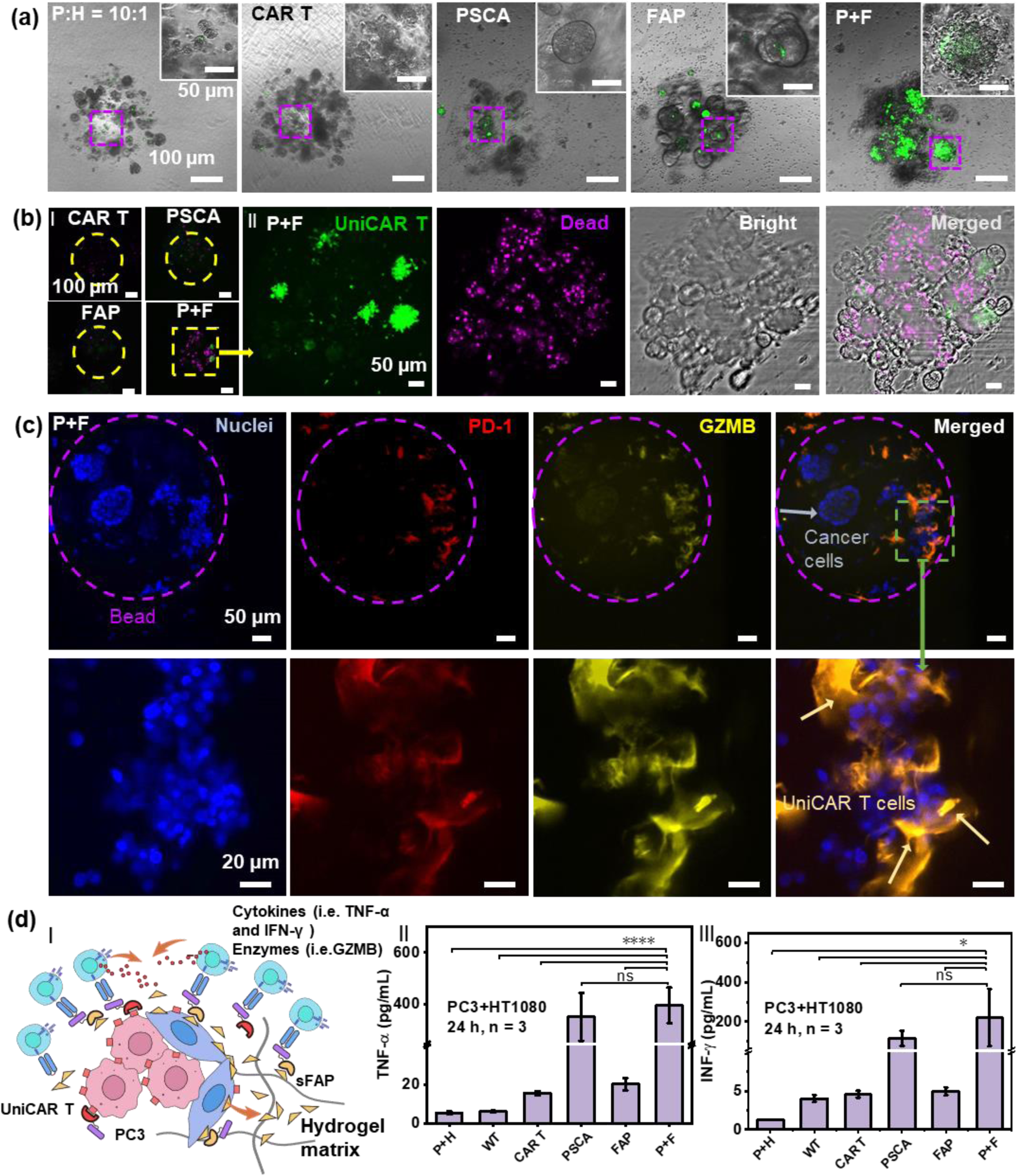
Therapeutic efficiency of T cells against P+H (10:1) spheroids. (a) Representative accumulation of UniCAR T cells (green) on P+H (10:1) spheroids in hydrogel beads after 48 h of culturing under different conditions. (b) Representative images of dead cell (magenta) staining treated by UniCAR T cells (green) after 48 h of culturing under different conditions. (c) Representative immunostaining of T cell markers (blue: nuclei; red: PD-1; yellow: Granzyme B (GZMB)). (d) (I) Illustration of possible scenarios for UniCAR T cells in PEGDA hydrogel beads; Concentrations of TNF-α (II) and IFN-γ (III) in the supernatant obtained after 24 h of culturing under different conditions. P+H: only P+H spheroids; WT: P+H spheroids with wild-type T cells; CAR T: P+H spheroids with UniCAR T cells; PSCA: P+H spheroids with UniCAR T cells and anti-PSCA TMs; FAP: P+H spheroids with UniCAR T cells and anti-FAP TMs; P+F: P+H spheroids with UniCAR T cells, anti-PSCA TMs, and anti-FAP TMs. p-values were calculated using the One-way ANOVA combined with Tukey’s multiple comparison test. Differences between experimental groups were considered significant when ns(0.1234), *p < 0.0332, **p < 0.0021, ***p < 0.0002, and ****p < 0.0001, n ≥ 3.

## 3. Discussion

To date, CAR T cell therapy demonstrates relatively low efficiency in the clinical treatment of solid tumors.^57^ The scientific community believes that one of the main causes is the immunosuppressive role of the tumor microenvironment, composed of the extracellular matrix and stromal cells, e.g., cancer-associated fibroblasts (CAFs) and mesenchymal stromal cells.^37, 58^ To improve therapeutic strategies, we need to acquire a better understanding of cancer progression and new knowledge about ‘unlocking the tumor microenvironment’ via bioengineering of tumor tissues, while taking into account the impact of TME. FAP, being overexpressed on CAF cells and present in soluble form (sFAP) in the extracellular matrix, is one of the crucial elements of TME that contributes to the remodeling of the extracellular matrix (ECM) and degradation of the links within the tumor tissue, thereby simplifying its potential metastatic activity.^34–36^ FAP was also known to influence signaling pathways, e.g., the ones controlled by nuclear factor-κB, which are relevant to cancer metastasis and cytokine-induced apoptosis.^54, 59^

In this study, we aimed to model the impact of FAP-expressing cells on the proliferation of 3D prostate cancer model in hydrogel microenvironments, and on the corresponding infiltration as well as immunotherapeutic efficiency of the redirected UniCAR T cells in the presence of TMs. For this purpose, we successfully prepared PC3-PSCA/HT1080 hFAP 3D multi-spheroid tumor models cultured for more than 20 days, incorporating tumor cells, artificial supporting matrix, and FAP positive cells co-cultured at different ratios using a microfluidic droplet generation and rapid UV polymerization to form PEGDA hydrogel beads. System parameters for the formation of PEGDA hydrogel beads, like UV intensity, monomer and photoinitiator concentrations, and flow rates, can be fine-tuned to adjust the important physical properties of the cell culture matrices, such as stiffness and permeability.

Interestingly, we observed a reduced size of the co-cultured PC3-PSCA and HT1080 hFAP spheroids (*ca.* 50–100 μm) compared to the PC3-PSCA mono-cultured spheroids (*ca.* 300 μm). This may be attributed to the effect sFAP has on the degradation of extracellular matrix components, such as collagen^29^. Immunohistochemistry staining confirmed the presence and distribution of the FAP in the matrix around the PC3-PSCA cells, depending on the co-culture ratio of the PC3-PSCA to HT1080 hFAP cells. In this respect, the obtained multi-spheroid model may resemble the initial and later stages of metastatic tumors in terms of cell distribution and FAP expression.^52^

Finally, these 3D multi-spheroid tumor models were also utilized to evaluate the infiltration of UniCAR T cells into the hydrogel matrix to reach the tumor regions and the efficiency of immunotherapy implemented using the UniCAR approach. Thanks to the easy *in situ* reprogramming of the UniCAR T cells, we demonstrated the benefits of simultaneous dual targeting of the tumor marker PSCA and the tumor microenvironment marker FAP, reflected in increased efficiency of cancer cell killing. UniCAR T cells successfully navigated through the physical barriers of the artificial PEGDA hydrogel matrix in the presence of anti-PSCA and anti-FAP TMs. The coordinated migration behavior of UniCAR T cells resulted in the selective accumulation on the surface of tumor spheroids, thereby enhancing the killing efficiency. These findings indicate that FAP acts as a physical or chemical shield for PSCA-positive cells, yet it remains a promising target for cancer therapy,^30, 60^ despite unclear mechanisms of soluble FAP formation.^50^ The possibility to detect cytokines and cytotoxic molecules showcases that our model facilitates cancer studies not only at the cellular level but also down to the molecular level.

Looking ahead, although artificial models cannot be a general replacement for natural organisms in all contexts of cancer research, high-throughput fabrication of reproducible miniaturized 3D cancer cell cultures still brings significant benefits. These models can open new avenues for reliable studies, accelerate research into complex cancer mechanisms, facilitate rapid *ex vivo* testing of therapies, and potentially reduce animal use. Further investigation in innovative material designs and integration of microfluidic systems can enable tunable bioreactors, allowing exploration of different facets of these systems, increasing their complexity, and broadening their potential applications. We are confident that this high-throughput approach for constructing complex tumor models is indispensable for transitioning from using defined cell lines to patient-derived samples, which can improve the representation of natural tumor heterogeneity, enhance the understanding of immunotherapeutic efficiency, and facilitate strategic planning of personalized therapies.

## 4. Methods and experiments

### 4.1 Materials

In this study, the following reagents were used without additional purification: poly(ethylene glycol) diacrylate (PEGDA, Mn 6000, Sigma-Aldrich), lithium phenyl-2,4,6-trimethylbenzoylphosphinate (LAP, Sigma-Aldrich), mineral oil (Sigma-Aldrich), Tween 20 (Sigma-Aldrich), phosphate-buffered saline (PBS, Sigma-Aldrich), Dulbecco’s phosphate-buffered saline (DPBS, Sigma-Aldrich), sucrose (Carl Roth), gelatin (Carl Roth), calcein AM and propidium iodide (PI) (VWR (Corning)), hematoxylin (Sigma-Aldrich), eosin B (Sigma-Aldrich), ROTIMount (Carl Roth), Roticlear (Carl Roth).

### 4.2 Preparation of PEGDA hydrogel beads and size measurement

PEGDA hydrogel beads were prepared at room temperature (20 ℃) in a water-in-oil system using our in-house built platform combining the T-junction-based microfluidic droplet generation with a setup for rapid UV polymerization, which was adapted from a previously reported droplet-based real-time monitoring system.^61–63^ The UV polymerization was carried out using a UV lamp (M365LP1, THORLABS) driven by constant electrical current sourced from a programmable DC power supply (SPD3303X-E, SIGLENT). A cell culture medium (RPMI STINO, described in section 4.5) supplemented with polymerization precursors (10% (w/v) PEGDA 6000 and 1% (w/v) LAP) was used as the water phase (WP). Mineral oil (Sigma-Aldrich) was used as the oil phase (OP). Liquids were stored in syringes and pumped into fluorinated ethylene propylene (FEP) tubing with an outside diameter of 1.6 mm and an inside diameter of 0.5 mm). The used flow rates were: q_WP_ = 5.5 μL‧min^−1^ and q_OP_ = 110 μL‧ min^−1^. Droplets were first generated in the polytetrafluoroethylene (PTFE) T-junction (inner diameter of 0.5 mm) and then quickly cross-linked by localized exposure to UV light (mean intensity: 0.23–0.35 W‧cm^−2^; exposure time: ∼2.55 s) through the FEP tubing. The UV light intensities were measured using a UV intensity meter coupled with a 365-nm sensor (diameter: 6.45 mm, Karl Suss, model 1000). The measured intensities correspond to the average intensities on the outside surface of the FEP tube. The experimental details and calculations related to UV irradiation are provided in the Supporting Information (SI Note and Figure S1).

Sizes of PEGDA hydrogel beads were determined from wide-field images obtained by an inverted microscope (Axio Observer, Zeiss). ImageJ analysis software was used to determine the particle size distribution. The distributions obtained from image analysis were fitted with a Gaussian distribution, giving the mean size and standard deviation (SD), as presented in the inset of Figure 1c. The polydispersity (*P*) expressed in percent was calculated by dividing the SD by the mean in the particle size distribution. The beads were considered as monodisperse for *P* < 0.1.

### 4.3 Physicochemical characterization of PEGDA hydrogel beads

Freshly prepared PEGDA hydrogel beads were initially rinsed with PBS-T (PBS containing 0.05% v/v Tween 20) three times. Following a single wash with DI water, the hydrogel beads were frozen at –20 ℃ overnight and then dried (Alpha 2-4 LSC plus, Christ) for 36 h. Scanning electron microscope (SEM, Phoenix XL) and Fourier transform infrared spectrometer (FTIR, Perkin Elmer, Inc.) were used to characterize the microstructure and chemical composition, respectively, as reported in our previous study.^12^

### 4.4 Mechanical stiffness test

Young’s moduli of PEGDA hydrogel beads were measured on a MicroTester G2 system (CellScale) in DPBS at 37 °C. Briefly, randomly selected PEGDA hydrogel beads were placed into the test chamber filled with DPBS. A 1 mm-by-1 mm plate was affixed to one end of a tungsten beam with a diameter of 0.1524 mm. Parallel plate compression was performed, which subjected the beads to 25% compression at a controlled rate of 30 μm‧s^−1^. The recorded force-displacement data was used to plot the stress-strain curves and calculate the Young’s moduli (E) as follows:

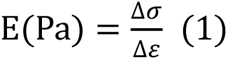

where *σ* refers to stress and *ε* refers to strain. Stress and strain were calculated according to the following equations:

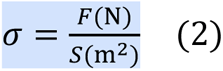

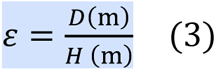

where *F* refers to force, *S* refers to the cross-sectional surface area of hydrogel beads, *D* refers to tip displacement, and *H* refers to the original height of hydrogel beads.

### 4.5 Cell culture

The prostate cancer cell line PC3 and the fibrosarcoma cell line HT1080 were obtained from the American Type Culture Collection (ATCC). The PC3 overexpressing PSCA (PC3-PSCA) and the HT1080 overexpressing human FAP (HT1080 hFAP) were prepared and confirmed by flow cytometry as described in previous reports.^64, 65^ PC3-PSCA cells were cultured in RPMI STINO medium: Roswell Park Memorial Institute (RPMI 1640 –Gluta, Gibco) supplemented with 10% fetal bovine serum (FBS, Biochrom), 1% streptomycin and penicillin (Biochrom), 1% MEM non-essential amino acid solution (Sigma-Aldrich), 1 mM sodium pyruvate solution (Sigma-Aldrich), and 2 mM Ala-Gin (Sigma-Aldrich). HT1080 hFAP cells were cultured in DMEM STINO medium: Dulbecco’s Modified Eagle Medium (DMEM + 4.5 g·L^−1^ D-Glucose, – Sodium Pyruvate, Gibco) supplemented with 10% fetal bovine serum (Biochrom), and 1% MEM non-essential amino acid solution (Sigma-Aldrich). Cells were incubated under 5% CO_2_ at 37 ℃. Cell culture medium was exchanged every 2–3 days.

### 4.6 Incorporation of cells into PEGDA hydrogel beads, culturing, and live/dead imaging A solution of RPMI STINO medium containing 10% (w/v) PEGDA 6000, 1% (w/v)

LAP, and predefined concentrations of cells was used as WP. Mono-cultured spheroids were generated using PC3-PSCA cells at a concentration of 1.2–1.5×10^7^ cells‧mL^−1^ and HT1080 hFAP cells at a concentration of 3.0–3.2×10^7^ cells‧mL^−1^. For co-cultured spheroids, a mixture of PC3-PSCA and HT1080 hFAP cells with concentration ratios of 10:1 and 2:1 was utilized to form a total concentration of 1.2–1.5×10^7^ cells‧mL^−1^. Mineral oil was employed as OP. The flow rates for WP and OP were the same as the ones described in section 4.2. The applied UV light intensity was 0.23 W‧cm^−2^. RPMI STINO medium was used to culture hydrogel beads containing cells, and the medium was refreshed every 2–3 days. Cell proliferation and spheroid growth were monitored using an optical microscope (Axio Observer, Zeiss). The average diameters of randomly selected spheroids were manually measured on micrographs using ImageJ (version: v1.54i). As for live/dead assay, spheroids in PEGDA hydrogel beads were firstly stained with calcein AM for 2 h (the first 10 days of culturing) or 3 h (after more than 10 days of culturing), and then with PI for 1 min. The solutions of staining chemicals were prepared following the instructions from the supplier (VWR (Corning)). Images were recorded using the confocal fluorescence microscope (Olympus IX83) coupled to an excitation dichroic mirror (DM) (DM405/488/543/635), emission DM (DM560), and band-pass filters (500–530 nm and 555–655 nm).

### 4.7 Isolation and genetic modification of T cells

Isolation of T cells and generation of UniCAR T cells were carried out as described in recent studies.^64, 65^ This study was approved by the local ethics committee of the Medical Faculty Carl Gustav Carus, Technical University Dresden (EK27022006). Briefly, T cells were isolated by using a human Pan T Cell Isolation Kit (Miltenyi Biotech GmbH, 130-096-535) from buffy coats provided by the German Red Cross (Dresden, Germany) after written consent of the voluntary donors. T cells without any modification were used as WT T cells. UniCAR T cells were generated by lentiviral transduction using a multiplicity of infections of 1–2. The proportion of UniCAR T cells was assessed via flow cytometry based on the co-translated Enhanced Green Fluorescent Protein (EGFP) marker protein expression prior to each experiment.

### 4.8 Production of TMs

The anti-PSCA-E5B9 and anti-FAP-E5B9 TMs were prepared as described previously.^64, 65^ Briefly, 3T3 cell lines expressing either anti-PSCA-E5B9 or anti-FAP-E5B9 TMs were produced by using lentiviral gene transfer. TMs were purified via His-Tag using nickel-nitrilotriacetic acid affinity chromatography (Qiagen GmbH). SDS-PAGE (Sodium dodecyl-sulfate polyacrylamide gel electrophoresis) and Western Blot were used to determine the concentration and purity of the TMs.

### 4.9 Immunostaining of spheroids

To observe the 3D structure of PC3-PSCA, HT1080 hFAP, PC3-PSCA and HT1080 hFAP (P+H) spheroids, the nuclei, PSCA, and FAP were targeted with Hoechst 33258 (ThermoFisher, 1:10), PSCA antibody (conjugated with Alexa Fluor 594, BIOSSUSA, 1:100), and hFAP antibody (R&D, 1:100), respectively. A secondary antibody Chicken anti-Mouse IgG (H+L) (conjugated with Alexa Fluor 647, Thermo Fisher Scientific, 1:50) was used to target hFAP antibody. To stain the spheroids directly in PEGDA hydrogel beads precultured for 7 days, the beads were transferred to 18 µ-slides and washed three times with PBS. Afterward, the PBS containing 4% paraformaldehyde was used to fix the spheroids for 10 min. Following the fixation step, the beads were incubated for 5 min in 10 mM EDTA to dissolve the gel matrix and release the spheroids. After permeabilization with 0.2% Triton X-100 for 10 min, the spheroids were blocked using 3% BSA solution in PBS for 2 h. Primary antibodies (both, anti-PSCA and anti-hFAP) were then added to incubate the spheroids at 4 °C overnight. After washing twice with tris-buffered saline containing 0.1% Tween 20 (TBS-T) and once with TBS, a secondary antibody was added and incubated for 1 h. Finally, the spheroids were incubated with Hoechst 33258 for 10 min. After washing, Prolong (ThermoFisher) antifade reagent was added to prevent fluorescence quenching. Images were recorded using the confocal fluorescence microscope (Olympus IX83) coupled to the excitation DM (DM405/488/543/635), emission DM (DM490/640), and band-pass filters (425– 475 nm, 555–625 nm, and 655–755 nm).

To stain the sections of spheroids, PEGDA hydrogel beads containing spheroids were embedded into a mixed gel which contains 20% sucrose and 7.5% gelatin. Sections of 40 µm thickness were then created using a microtome cryostat under 22 ℃ (CM1850, Leica). After drying at 37 °C for 20 min, the sections were fixed with 4% paraformaldehyde for 10 min. The mixed gel can be washed off by using TBS-T buffer at 50–60 °C for approximately 10 min. The staining process for sections follows the same method as bead staining described previously, except for the primary antibody incubation parameters (2 h at room temperature).

### 4.10 T cell counting and migration tracking

PEGDA hydrogel beads with cancer spheroids were first prepared and cultured in a 6-well plate at 37 °C under 5% CO_2_ for 7 days. Then, for each tracking experiment, a single bead was transferred to one well of an 18-well µ-Slide (ibidi, uncoated). This step was followed by adding 50 μL of cell culture medium with WT T cells, UniCAR T cells, UniCAR T cells + anti-PSCA-E5B9 TMs, UniCAR T cells + anti-FAP-E5B9 TMs, or UniCAR T cells + anti-PSCA-E5B9 TMs + anti-FAP-E5B9 TMs, separately. Time-lapse images were captured using an optical microscope (Axio Observer, Zeiss) and monitored in ZEN3.3 software. Images were taken every 30 s over a time span of 1 h. Experiments were replicated at least three times under the same conditions. A circular region of interest with a diameter of 450 μm was manually selected to track the T cells using ImageJ (version: v1.54i). Then, the TrackMate 7^66^ (ImageJ plugin) was used to track the T cells moving inside and outside of the circular region. T cells were counted within a ring area with an inner diameter of 450 μm and an outer diameter of 800 μm. Hessian detector and LAP tracker were selected to accurately identify T cell positions in the captured images. The detection threshold was manually adjusted depending on the individual images.

### 4.11 T cell treatment experiments and killing efficiency

PEGDA hydrogel beads with cancer spheroids were first prepared and cultured in a 6-well-plate at 37 °C under 5% CO_2_ for 7 days. Then the beads were transferred to 96-well plates for T cell co-culturing incubation. Five beads with PC3-PSCA spheroids, HT1080 hFAP, or P+H spheroids were placed in each well. 200 μL of cell culture medium with WT T cells, UniCAR T cells, UniCAR T cells + anti-PSCA-E5B9 TMs, UniCAR T cells + anti-FAP-E5B9 TMs, or UniCAR T cells + anti-PSCA-E5B9 TMs + anti-FAP-E5B9 TMs was added to the wells. The concentration of T cells was 0.5×10^6^ cells·mL^−1^, and the concentration of TMs was 5 pmol·mL^−1^.

Killing efficiency was determined using PI staining (as described in section 4.6). After staining, the spheroids were imaged with the confocal fluorescence microscope. For the determination of TNF-α and INF-γ concentrations, cell supernatants were collected for enzyme-linked immunosorbent assay (ELISA) after co-culturing for 24 h. The procedures outlined in the protocols provided with the ELISA kits purchased from Thermo Fisher Scientific Inc. were followed.

T cell immune response markers PD-1 and Granzyme B (GZMB) were detected via immunostaining following the protocol for cell staining in hydrogel beads (refer to section 4.9 for details). Recombinant PD-1 antibody (conjugated with CF 488, antibodies-online, 2 μg·mL^−1^) and GZMB antibody (conjugated with Alexa Fluor 555, antibodies-online, 1:100) served as primary antibodies. Hoechst 33258 served as the contrast agent. Images were recorded using the confocal fluorescence microscope (Olympus IX83) coupled to the excitation DM (DM405/488/543/635), emission DM (DM490/560), and band-pass filters: 425–475 nm, 500–530 nm, and 555–625 nm.

### 4.12 Statistical analysis

Data are reported as mean ± SD. p-values were calculated using the One-way ANOVA combined with Tukey’s or Dunnett’s multiple comparison test using GraphPad Prism 9 depending on the analyzed dataset. Differences between experimental groups were considered as significant when ns(0.1234), *p < 0.0332, **p < 0.0021, ***p < 0.0002, and ****p < 0.0001, n ≥ 3.

## Supporting Information

Supporting Information is available from……or from the author.

## Supporting information

preprint-3D Microphysiological Tumor Model for Dual-Targeting CAR T Cell Immunotherapy

## Acknowledgements

We would like to thank Aline Morgenegg, Johanna Wodtke, Xinne Zhao, Ph.D., Trang Anh Nguyen Le, and Diana Isabel Sandoval Bojorquez from the Institute of Radiopharmaceutical Cancer Research. We also thank for the support from the China Scholarship Council (No. 202006630001), the LiSyM Cancer phase I joint collaborative project DEEP-HCC (Federal Ministry of Education and Research; No. 031L0258B), Deutsche Forschungsgemeinschaft (DFG, Nr.BA4986/8−1), GRK 2767, and Helmholtz Initiative and Networking Fund in the project ‘Mhelthera’ (project ID: InterLabs-0031). Furthermore, LB acknowledges the financial support of the European Research Council (ERC) in the ERC-Consolidator Grant (ImmunoChip, 101045415).

