## Supplementary material for "3D Microphysiological Tumor Model for Dual-Targeting CAR T Cell Immunotherapy": preprint-3D Microphysiological Tumor Model for Dual-Targeting CAR T Cell Immunotherapy

### Supporting Information

#### SI Note: UV irradiation experiments and calculations

*UV intensity*. Due to the limited measurement range of the UV light sensor, we measured the intensities of emitted UV light when the lamp was driven by the DC electric current in the range from 0.05 to 0.55 A, and then calculated the linear fit to extrapolate the UV light intensities obtained under the current drive in the range from 1 to 1.5 A (defined as measured intensity, see Figure S1b). The illustration of the measurement setup geometry is shown in Figure S1c. The peak intensity (directly under the UV LED) and the mean intensity contributing to polymerization at the level of the FEP tube top surface were calculated by using MATLAB (R2022b) based on the measured intensity and the spatial radiation distribution data from the M365LP1-SpecSheet provided by the supplier (Table S1). The distance between the UV LED and the top surface of the FEP tube was measured using a digital micrometer (Schut).

Table S1. UV light intensity produced for different supplied electric currents.

| Electric current (A) | Intensity ($W \cdot cm^{-2}$) | | |
|---|---|---|---|
| | Measured | Peak | Mean |
| 1.00 | 0.36 | 0.48 | 0.23 |
| 1.25 | 0.46 | 0.62 | 0.29 |
| 1.50 | 0.55 | 0.74 | 0.35 |

*Exposure time calculation.* The injection flow rates for cell culture medium and mineral oil were $q_{WP} = 5.5\ \mu L \cdot min^{-1}$ and $q_{OP} = 110\ \mu L \cdot min^{-1}$, respectively. The total amount of liquid flowing through the tube per min ($V_{min}$) was 115.5 $mm^3$. The inside diameter of the tube is 0.5 mm, and thus, the inner cross-sectional surface area ($S_{in}$) is $0.25^2 \cdot \pi\ mm^2$ ($\pi \approx 3.14$). The length of liquid flowing through the tube per minute ($L_{min}$) was $V_{min}/S_{in}$ = 588.24 mm. The length of the tube placed within the irradiated area ($L_{irr}$) was 25 mm. Hence, the exposure time (equivalent to the time required for the liquid to pass through the irradiated area) can be calculated as $(L_{irr}/L_{min}) \cdot 60\ s \approx 2.55\ s$.

#### SII Note: FTIR analysis

The band at 2888 cm$^{-1}$ can be assigned to the asymmetric stretching of $CH_2$. The bands at 1726 cm$^{-1}$ and 1098 cm$^{-1}$ correspond to the symmetric stretching of C=O and stretching vibrations of C–O, respectively. The bands at 1645 cm$^{-1}$, 961 cm$^{-1}$, and 842 cm$^{-1}$ are assigned to the vibrations of the aliphatic double bond C=C, out-of-plane symmetric stretching of $CH_2$=CH, and symmetric stretching of $CH_2$=CH, respectively.[13, 38]

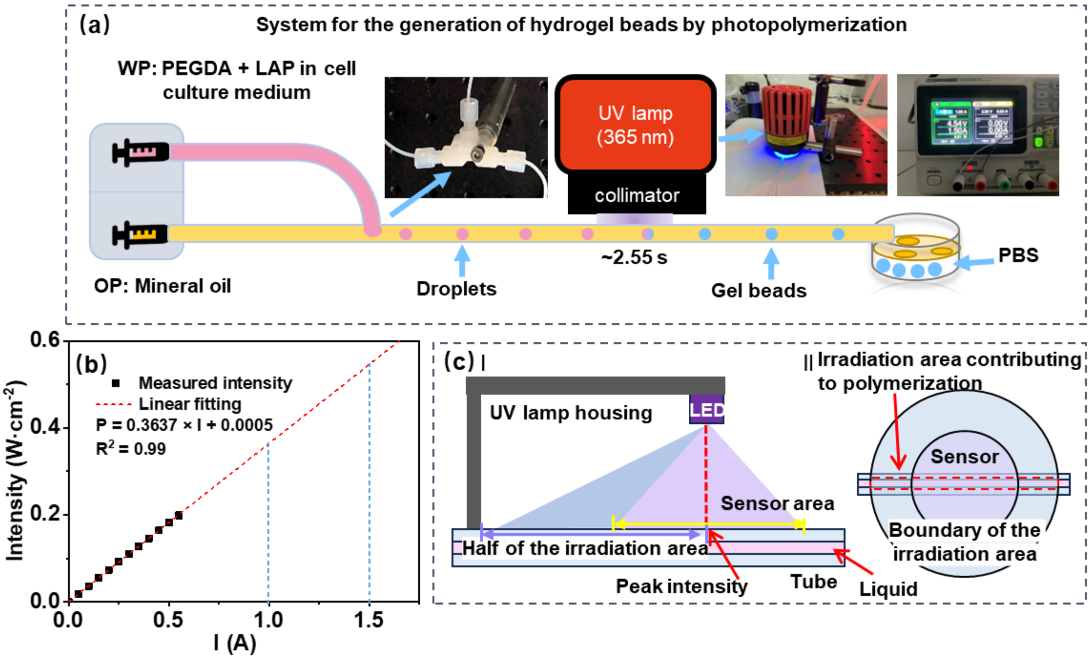


Figure S1. (a) Illustration and photographs of PEGDA hydrogel beads generation platform. (b) Measured UV intensity versus supplied direct current (DC) and the corresponding linear fit. (c) Illustration depicting the geometry of UV intensity measurements and the irradiated areas, shown from side view (I) and top view (II). WP: water phase; OP: oil phase.

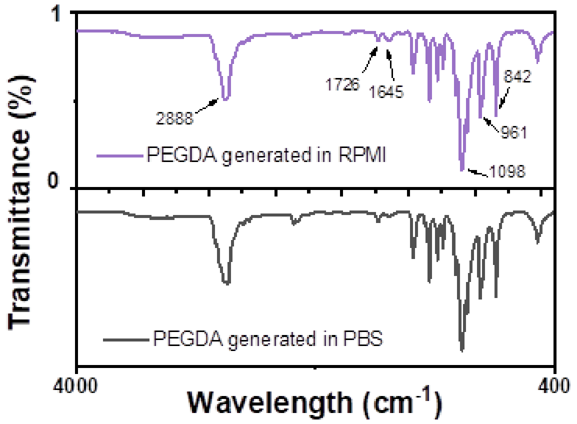


Figure S2. FTIR spectra of PEGDA hydrogel beads generated in RPMI-STINO cell culture medium and PBS.

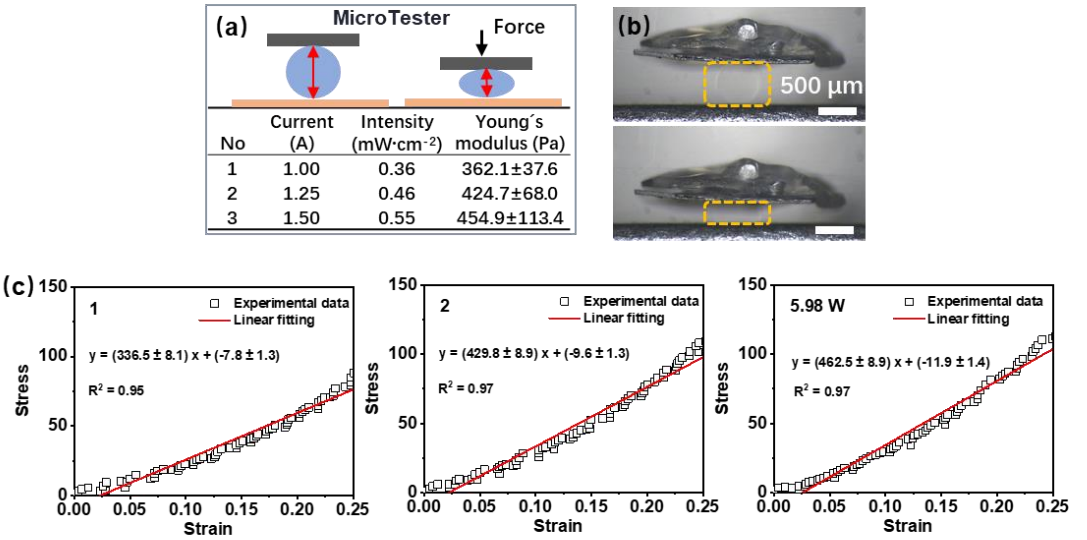


Figure S3. Stiffness of PEGDA hydrogel beads generated under different UV light intensities. (a) Illustration of the stiffness testing process and summary of stiffness values for hydrogel beads prepared under different UV light intensities. (b) Representative images showcasing the testing process. (c) Representative stress-strain curves with corresponding linear fits used to determine the stiffness of PEGDA hydrogel beads (n = 5).

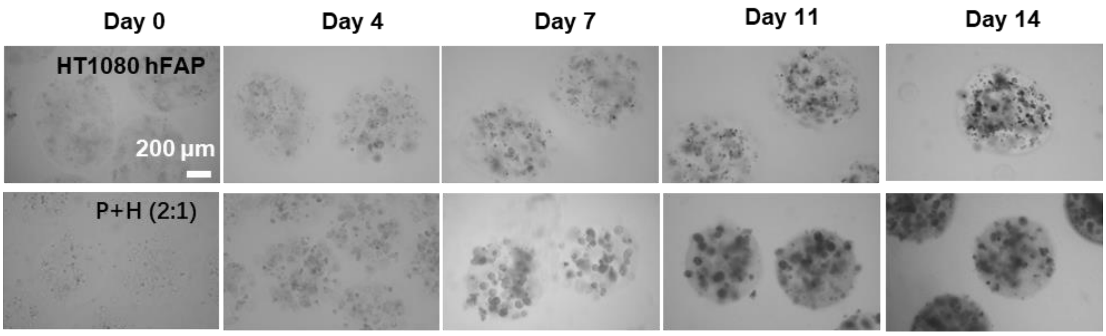


Figure S4. Representative optical micrographs illustrating the processes of spheroid formation and proliferation in PEGDA hydrogel beads loaded with HT1080 cells or PC3 and HT1080 cells (P:H = 2:1).

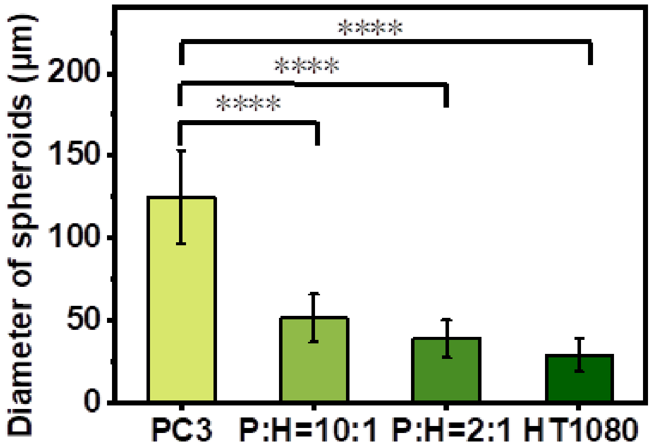


Figure S5. Average diameter of different spheroids measured on day 14, n = 100. p-values were calculated using the One-way ANOVA combined with Dunnett's multiple comparison test. Differences between experimental groups were considered as significant when *$p < 0.0332$, **$p < 0.0021$, ***$p < 0.0002$, and ****$p < 0.0001$, n ≥ 3.

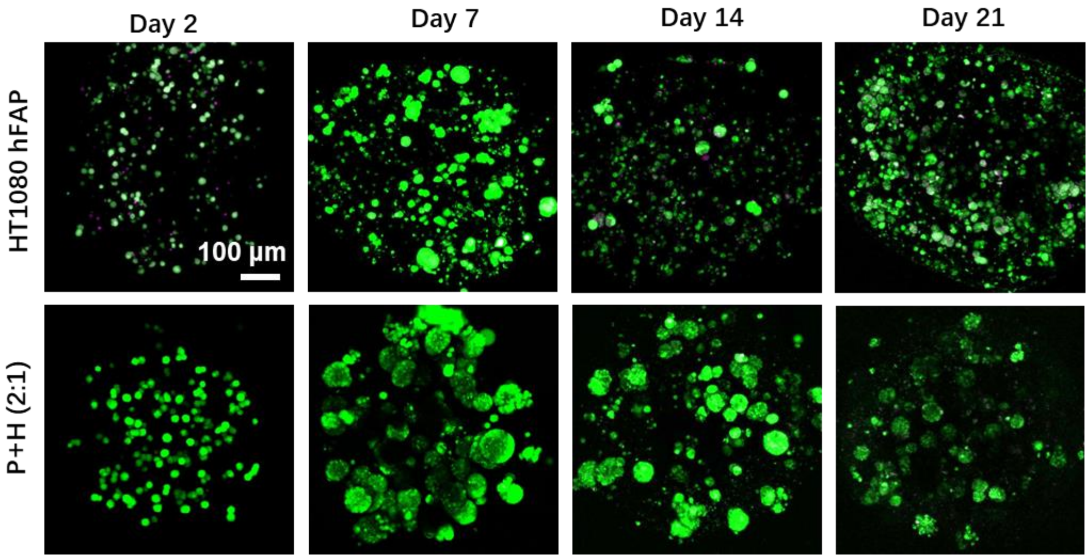


Figure S6. Representative fluorescent micrographs of live/dead (green/magenta) staining for P+H (2:1) and HT1080 hFAP spheroids.

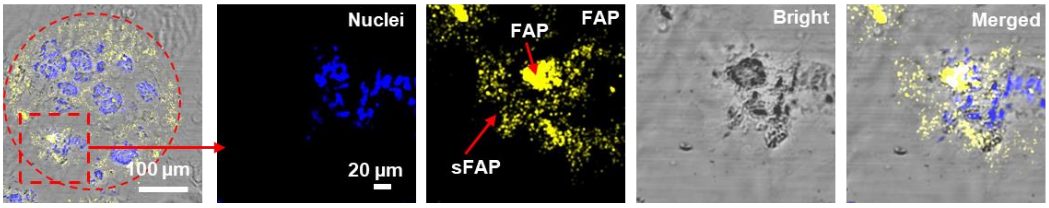


Figure S7. Immunostaining images of spheroid sections illustrating the distribution of FAP and sFAP.

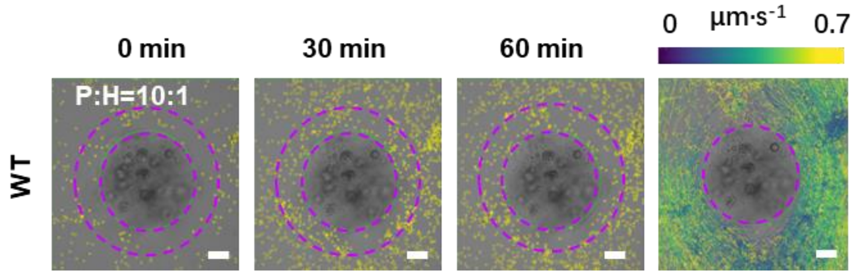


Figure S8. Representative images of wild-type (WT) T cell distribution and tracking of cell paths over 1 h.

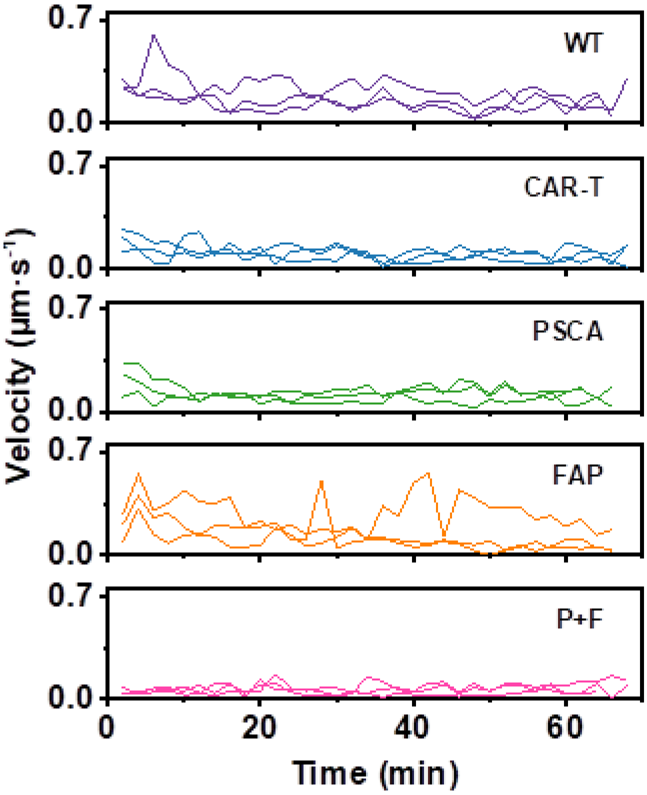


Figure S9. Representative velocities as a function of time for T cells under different microenvironmental conditions. **WT**: P+H (10:1) spheroids with wild-type T cells; **CAR T**: P+H spheroids with universal CAR T cells; **PSCA**: P+H spheroids with universal CAR T cells and anti-PSCA TMs; **FAP**: P+H spheroids with universal CAR T cells and anti-FAP TMs; **P+F**: P+H spheroids with universal CAR T cells, anti-PSCA TMs, and anti-FAP TMs.

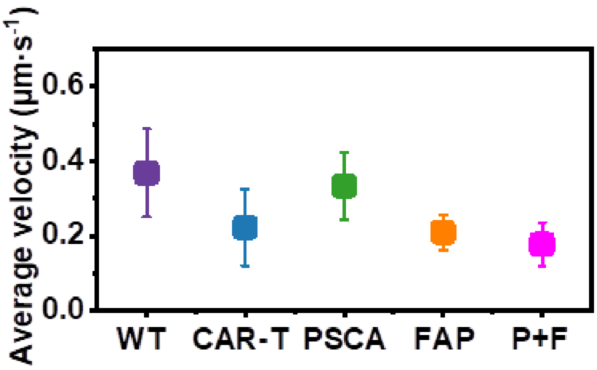


Figure S10. Average velocity of T-cell motion under different microenvironmental conditions calculated from Figure 3a(II). **WT**: P+H (10:1) spheroids with wild-type T cells; **CAR T**: P+H spheroids with universal CAR T cells; **PSCA**: P+H spheroids with universal CAR T cells and anti-PSCA TMs; **FAP**: P+H spheroids with universal CAR T cells and anti-FAP TMs; **P+F**: P+H spheroids with universal CAR T cells, anti-PSCA TMs, and anti-FAP TMs.

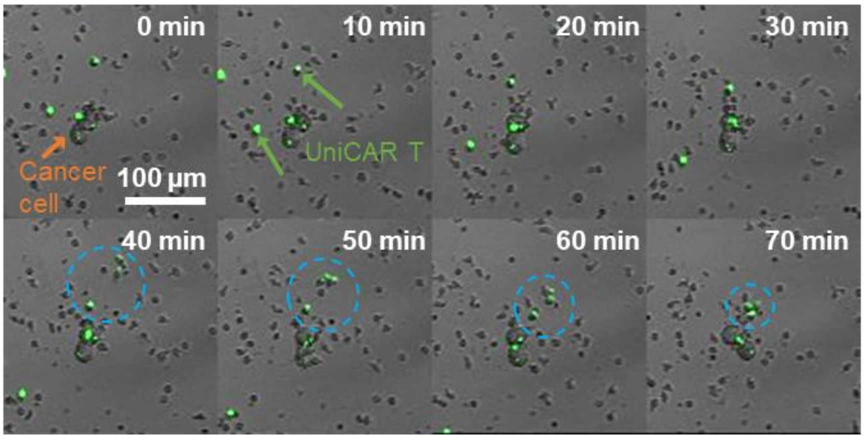


Figure S11. A representative example of universal CAR T cell accumulation around a cancer cell in plate culture in the presence of anti-PSCA and anti-FAP TMs.

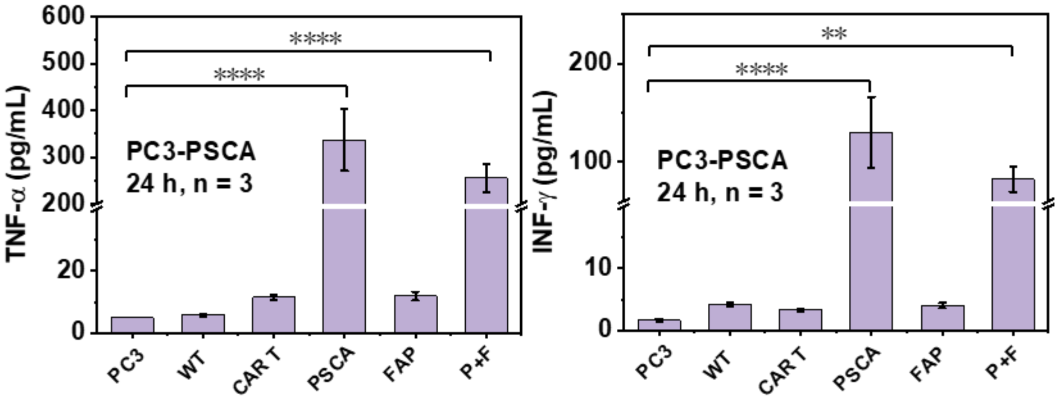


Figure S12. Concentrations of TNF-α and IFN-γ in the supernatant after 24 h of culturing under different conditions. PC3: only PC3-PSCA spheroids; WT: P+H spheroids with wild-type T cells; CAR T: PC3-PSCA spheroids with universal CAR T cells; PSCA: PC3-PSCA spheroids with universal CAR T cells and anti-PSCA TMs; FAP: PC3-PSCA spheroids with universal CAR T cells and anti-FAP TMs; P+F: PC3-PSCA spheroids with universal CAR T cells, anti-PSCA TMs, and anti-FAP TMs. p-values were calculated using the One-way ANOVA combined with Tukey's multiple comparison test. Differences between experimental groups were considered as significant when *p < 0.0332, **p < 0.0021, ***p < 0.0002, and ****p < 0.0001, n ≥ 3.

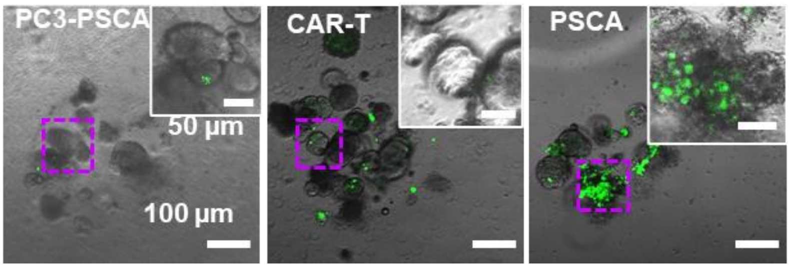


Figure S13. Universal CAR T cell (green) accumulation on PC3-PSCA spheroids after 48 h of culturing under different conditions. PC3-PSCA: only PC3-PSCA spheroids; CAR T: PC3-PSCA spheroids with universal CAR T cells; PSCA: PC3-PSCA spheroids with universal CAR T cells and anti-PSCA TMs.

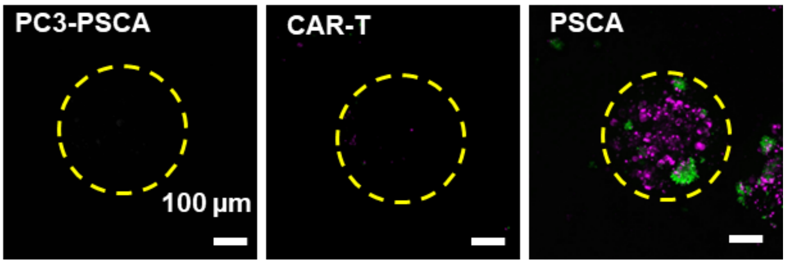


Figure S14. Dead cell (magenta) staining of PC3-PSCA spheroids after 48 h of culturing under different conditions. Green: universal CAR T cells. PC3-PSCA: only PC3-PSCA spheroids; CAR T: PC3-PSCA spheroids with universal CAR T cells; PSCA: PC3-PSCA spheroids with universal CAR T cells and anti-PSCA TMs.

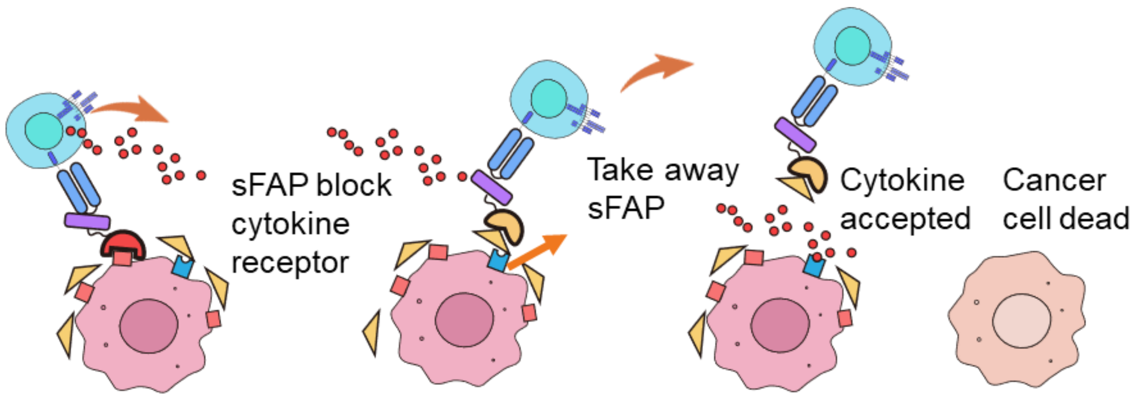


Figure S15. Hypothesis of the mechanism of the PSCA and FAP dual-targeting system.

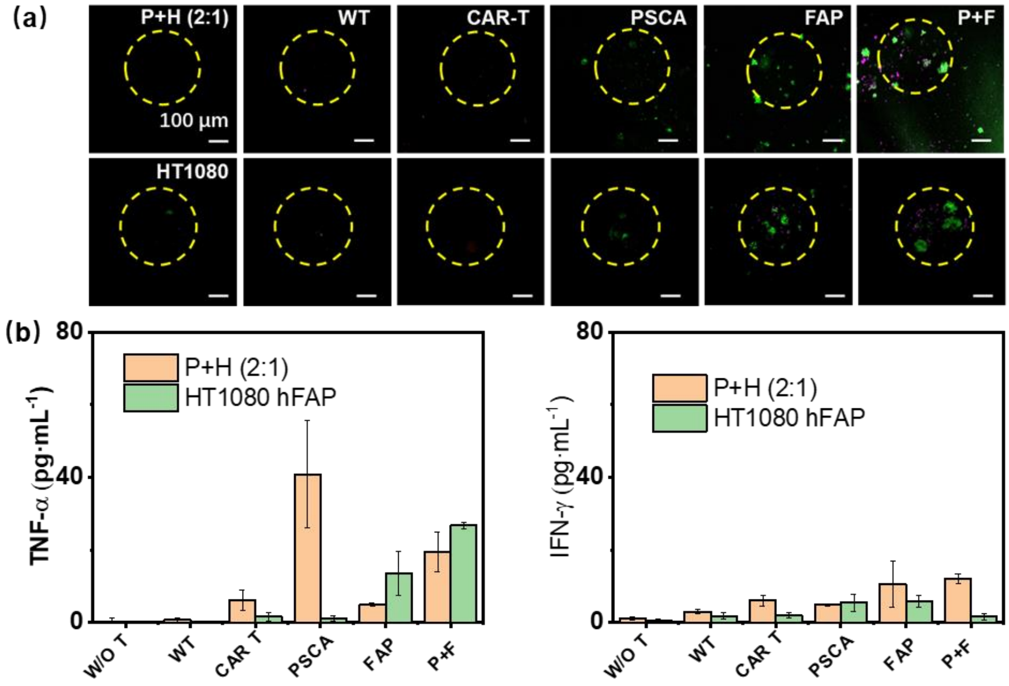


Figure S16. Tumor cell killing efficiency of universal CAR T cells in P+H (2:1) and HT1080 hFAP 3D *in vitro* tumor models. (a) Dead cell (magenta) staining of P+H (2:1) and HT1080 hFAP spheroids after 48 h of culturing. (b) Concentrations of TNF-α and IFN-γ in the supernatant after 24 h of culturing under different conditions. Green: universal CAR T cells. PC3-PSCA: only spheroids; CAR T: spheroids with universal CAR T cells; PSCA: spheroids with universal CAR T cells and anti-PSCA TMs. FAP: spheroids with universal CAR T cells and anti-FAP TMs; P+F: spheroids with universal CAR T cells, anti-PSCA TMs, and anti-FAP TMs.
